# Simultaneous Profiling of Host Expression and Microbial Abundance by Spatial Meta-Transcriptome Sequencing

**DOI:** 10.1101/2022.08.04.502882

**Authors:** Lin Lyu, Xue Li, Ru Feng, Xin Zhou, Tuhin K. Guha, Xiaofei Yu, Guo Qiang Chen, Yufeng Yao, Bing Su, Duowu Zou, Michael P. Snyder, Lei Chen

## Abstract

We developed an analysis pipeline that can extract microbial sequences from Spatial Transcriptomic (ST) data and assign taxonomic labels, generating a spatial microbial abundance matrix in addition to the default host expression matrix, enabling simultaneous analysis of host expression and microbial distribution. We called the pipeline Spatial Meta-transcriptome (SMT) and applied it on both human and murine intestinal sections and validated the spatial microbial abundance information with alternative assays. Biological insights were gained from this novel data that demonstrated host-microbe interaction at various spatial scales. Finally, we tested experimental modification that can increase microbial capture while preserving host spatial expression quality, and by use of positive controls, quantitatively demonstrated the capture efficiency and recall of our methods. This proof of concept work demonstrates the feasibility of Spatial Meta-transcriptomic analysis, and paves the way for further experimental optimization and application.

## Introduction

Spatial transcriptomic (ST) sequencing has revolutionized the biological research field. This class of technologies combine the strength of two pillars of modern biological research, sequencing and imaging. One form of ST mapping works by capturing the messenger RNA from a permeabilized tissue slice, and label these RNA molecules with 2D spatial barcodes (Stahl et al. 2016; Salmen et al. 2018). Generally this has been applied to complex human tissues such as the brain and tumors (Berglund et al. 2018; Thrane et al. 2018; Asp et al. 2019; Maniatis et al. 2019; Moncada et al. 2020; Fawkner-Corbett et al. 2021; Maynard et al. 2021). In the intestines and other organs, microbes live alongside or within close proximity to host cells and meta-genomic sequencing has long been employed to study the complex microbial composition on various host body sites. These microbiome studies have greatly improved our understanding of the human biology and microbes have been found in novel places (Nejman et al. 2020; Poore et al. 2020) and showed intriguing dynamics across solid tissue (Ha et al. 2020). However, they generally lacked spatial resolution and attempts at addressing this limitation is just emerging (Duncan et al. 2020; Shi et al. 2020). Inspired by a recent study about COVID19 where virus sequences were recovered from host single cell transcriptomic data and analyzed alongside host data (Bost et al. 2020), we tested if it is possible to capture microbial sequences in spatial transcriptomic sequencing and demonstrated that validated microbial information can be obtained that can shed new light on host-microbe interactions.

## Results

### SMT extracts microbial abundances alongside host expression

To evaluate the microbial sequence content from spatial transcriptome data, we performed spatial transcriptomic sequencing, using the Visium kit from 10X Genomics™, on both human and murine intestinal samples, where the presence of microbes is well-known. The human samples included dissected colon samples from two colorectal cancer patients. For each patient, samples from the tumor site and histologically normal site distant from the tumor were embedded together and put in the same capture window and two replicate windows were processed. The murine samples came from both small intestine and large intestine, and cross-sectional slices from 3 mice were included in a single window (Fig. 1A, Fig S1C). The murine samples were rinsed so as to remove fecal material while preserving the mucus layer. The human samples were rinsed at the operation room before transported to downstream processing. Each slice was sequenced to at least 85k reads per covered spots, and the captured RNAs were on average 11k per spot, similar to that of published results (Stahl et al. 2016).

**Fig. 1.**
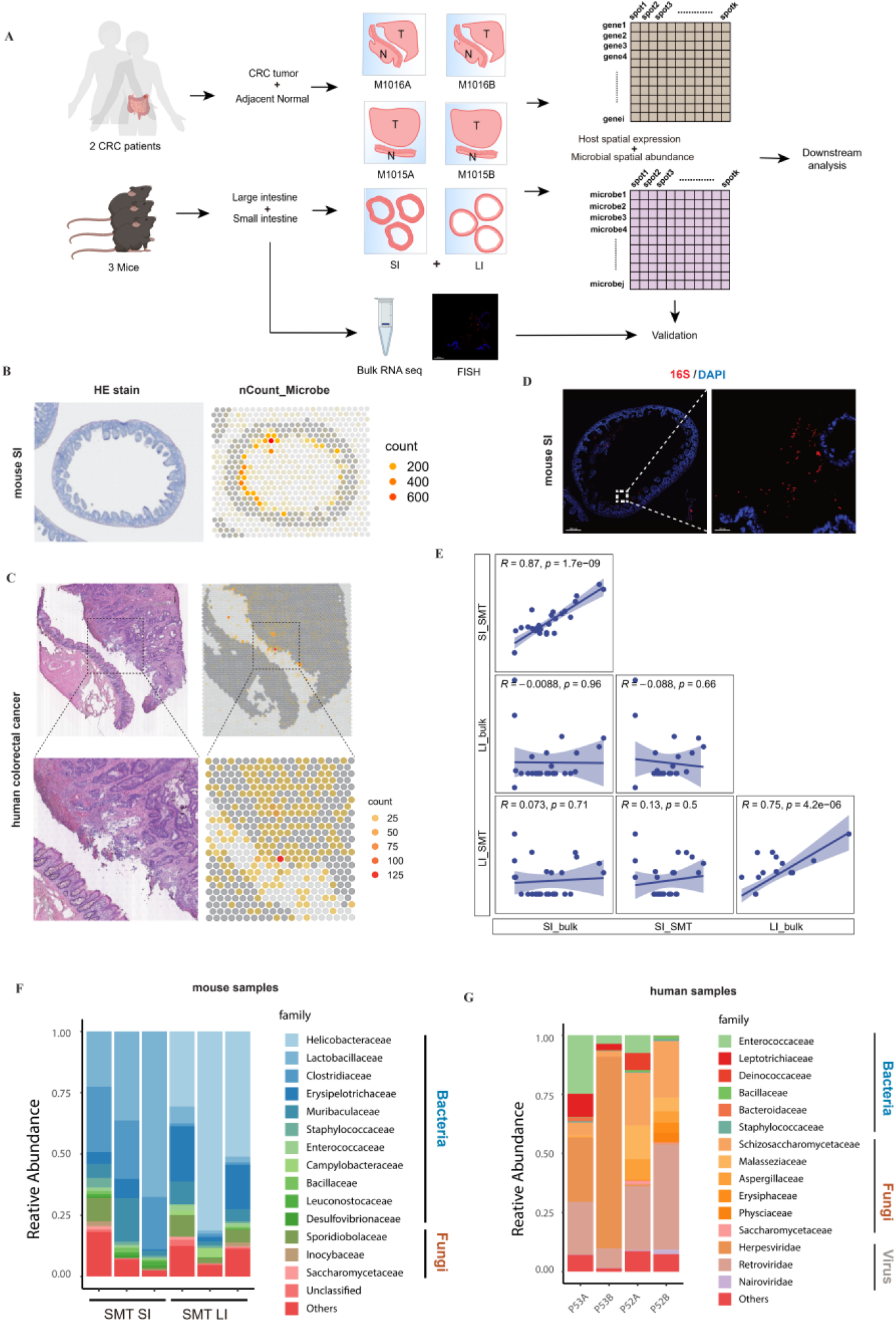
Spatial meta-transcriptomics of human and murine intestines. **(A)** A schema of the SMT framework and design of the first part of study, note that for CRC, tumor and adjacent normal tissue were embedded in the same window and designated T and N respectively. **(B-C)** H&E staining of the tissue sections (left) and corresponding spatial feature plot showing microbial sequence captured per spot captured (right) for murine small intestine (B) and a human colorectal cancer patient tissue, from patient M1016, with blown up (C). Representatives were shown and additional can be found in the Fig. S1. **(D)** FISH result for mouse small intestine section stained for bacteria 16S. Scale bars: 300μm (left), 20 μm (right). **(E)** Microbial abundance correlation between SMT generated pseudo-bulk measurements (x-axis) and bulk RNA-Seq measurements (y-axis), Spearman’s correlation co-efficiencies and p values were shown. **(F, G)** Stacked bar plot showing microbial abundances for mouse (F) and human (G) samples at family level, only the top ones were shown.

**Fig. S1. Related to Fig 1.**
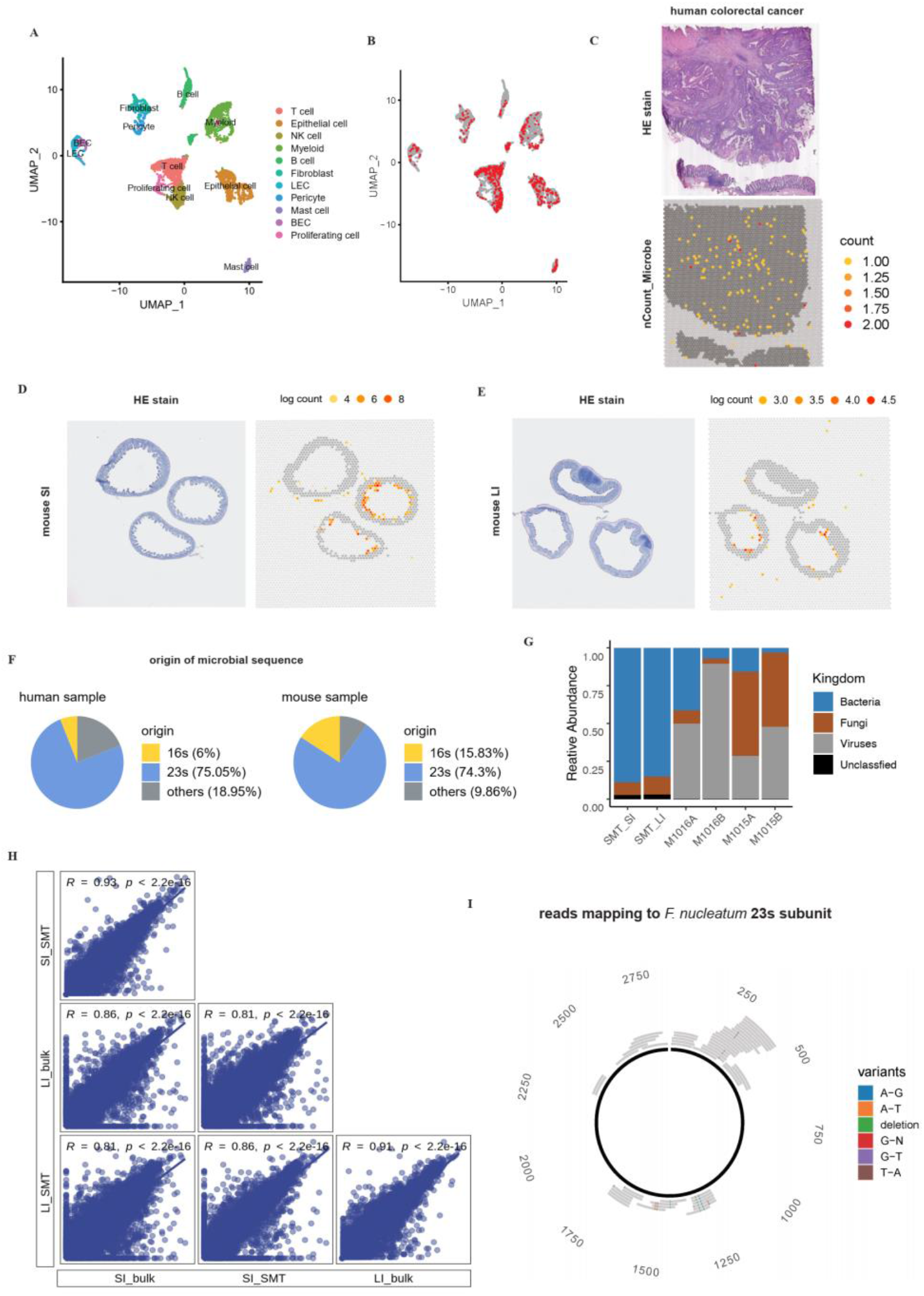
**(A)** TSNE plot showing single cell sequencing of an unsorted CRC sample, clustered into 11 major cell types. **(B)** Cells in (A) were highlighted in red if they were enriched in microbial sequences (>0.02%). **(C)** H&E stain (top) and microbial UMI count (bottom) for patient M1015. **(D-E)** H&E staining of the tissue section (left) and corresponding log scaled microbial sequence count per spot captured by SMT (right) for mouse small intestine (D) and large intestine (E). Note that 3 samples were placed in each window. **(F)** Pie chart showing origin of microbial sequence captured, for human samples (left) and murine samples (right). **(G)** Microbial composition cross human and murine samples at kingdom level. **(H)** Correlation for host genes between SMT pseudo-bulk measurements (x-axis) and bulk RNA-Seq measurements (y-axis). **(I)** Representative read alignment to the *Fusobacterium* 23S gene, for M1016A.

We developed an analysis pipeline to extract microbial information from the ST data. In short, reads unmapped to the host genome were first screened to remove low complexity reads and then aligned against the NCBI NT database using Blast and subsequently assigned taxonomic IDs. When there are conflicting assignments for reads belonging to the same Unique Molecular Identifier (UMI) group, a strategy combining majority voting and most recent common ancestor was used to resolve it. The end result is a spatially resolved microbial abundance matrix alongside the spatial host gene expression matrix generated via the standard ST analysis. We compared our procedure with Kraken 2 (Wood et al. 2019), a tool commonly used in taxonomy assignment for meta-genomics reads, and found our tailored procedure to produce less dubious calls (methods, Table S1).

**Table. S1.** SMTA & kraken2 comparison.

|  | Kraken2 | SMTA |
| --- | --- | --- |
| Microbial reads called<sup>a</sup> | 63493 | 38977 |
| Microbial UMI called | 8593 | 1453 |
| Read/UMI<sup>b</sup> | 7.3 | 26.8 |

a. Kraken2 and SMTA used same dataset, and the total number of input read is 3233863.
b. Used as a measure of quality, overall read/UMI is $717337698/27998444 = 25.6$ .

The percentage of microbial sequences in a dataset vary across different samples, ranging from 10^−6^ in a human sample to 10^−3^ in murine ones. The Visium kit uses poly T to capture messenger RNA and in theory microbial sequences without poly A tail would be depleted. While this was indeed the case, some random priming did occur and sequences without long stretches of poly A were still captured. This is in line with reports from the RNA-velocity work (La Manno et al. 2018), where immature messenger RNAs, which lack poly A tails, account for about 25% of the typical data from 10X Genomics Chromium single cell platform, which also uses poly A capture. Furthermore, ribosomal RNA is the most abundant RNA species in a cell, accounting for more than 80% of total RNA. Ribosomal RNA is also the molecular marker of choice for nucleated microbial taxonomy. Accordingly, the majority of the microbial sequences recovered in SMT were from ribosomal RNAs (Fig. S1F). In addition, most microbial species, except airborne fungi ones, exhibited clear spatial patterns (agglomerated microbial signal shown in Fig. 1B, C). For these reasons, the microbial signals obtained from our SMT analysis is unlikely to be merely artifacts. For comparison, we applied our microbial sequence extraction method on single cell sequencing data derived from un-sorted intestinal samples and the microbial signal thus produced was spread relatively evenly spread across different cell types and likely represent noise or contaminants (Fig S1A, B). This may be due to the fact that only intra-cellular bacteria can be captured in cleaned single cell suspension and these bacteria are rare and that the gentle treatment used inside a droplet is ineffective for lysing them.

### Evaluation of contamination level and biases in SMT

To further evaluate the extent of contaminating microbial sequences during experiments versus true organ resident microbial signals. We plotted the microbial abundance across all spatial spots, regardless of whether it’s covered by tissue or not (Fig 1B, C). For both human and mouse, the luminal side of the intestines harbor more microbial signal as is expected. The human tumor tissue also showed a higher abundance of microbes and also more microbes penetrated deeper into the tissue, as would be expected from the compromised barrier function in tumors and the their folded overgrowth. In addition, the pattern of microbial distribution is similar between replicates, both spatially and in terms of taxa composition (Fig 1F, Fig S1D). These clear and expected patterns validated our SMT methods as noise or contaminants would show up relatively randomly across the capture window.

To compare results with alternative methods, for mice small intestine, we visualized neighboring section of the embedded tissue by Florescent In Situ Hybridization (FISH) with the bacterial probe EUB338, targeting 16S gene (Fig 1D). The FISH result was in agreement with SMT, showing bacteria presence concentrated in the intestinal lumen side. Furthermore, each window contained samples from 3 mice and the bacteria loads of these samples differ. This difference could be bona fide biological difference, combined with variance in sample processing, or slight difference in sample origin, i.e. more distal or proximal. Indeed, the order of the bacteria loads were observed to be consistent in proximate sections used for FISH and SMT, as were expected. All these observations lend credibility to SMT.

Next, we evaluated the biases in the microbial signals from SMT. During the standard Visium ST process, the host tissue sections are permeabilized to release its RNA and this process is far from ideal for microbial sequence capture. This brought up the question if certain microbial species would be preferentially analyzed and represent a biased picture of the true microbiome (some biases is always expected). We were also concerned that if the capture efficiency of microbial sequences were too low there would be a great fluctuation in the results and they would be unreliable. To address these issues, for the embedded murine samples, we extracted RNAs from neighboring sections and performed total RNA sequencing, in which poly A based enrichment was not used and only host ribosomal RNA was depleted. Similar to our SMT pipeline, we processed the bulk total RNA-Seq data to generate microbial abundances, one abundance profile for one window (containing samples from 3 mice). To compare with bulk sequencing data, we then combined the SMT’s per spot abundance profiles for one window into a single pseudo-bulk profile and evaluated both the correlation of microbial abundances at the family level (Fig. 1E) as well as that for host genes (Fig. S1H). The Spearman’s correlation of the microbial abundances are positive and significant between bulk and pseudo-bulk for the same tissue and insignificant otherwise, again validating the SMT approach. The pattern is different for host genes, as the correlation are all significant and positive due to similarity in overall expression. Lastly, the microbial composition based on SMT were shown to be mostly of intestinal origin based on literature, and known common contaminants and artifacts were present but at low levels (Fig 1F, G) (Ha et al. 2020; Dohlman et al. 2021).

### The localized host cell response to Cytomegalovirus infection as captured by SMT

An advantage of SMT over 16S sequencing is that by using RNA-Seq, SMT captures fungi and viral sequences as well as bacteria ones. Zoonotic viruses generally have mRNA that resemble their hosts and interestingly we can identify one such virus infection case in great spatial detail in one of the CRC patient. In both replicate windows of patient M1016, viral sequences reaching as high as 3.4% were observed in some locations at the tumor site. This virus, identified as *Cytomegalovirus*, is known to infect fibroblasts, and a deconvolution of the infected spots, which were 55μm in diameter and usually contained many cells, showed them to be mostly composed of fibroblast (Fig. 2A). We next investigated the immediate consequence of viral infection on these fibroblast by looking at their expression. To this end, infected spots were compared to un-infected fibroblast-rich spots. Interestingly, among the differentially expressed genes, interferon related genes are not found, potentially due to the immune-compromised nature of the tumor micro-environment. Still, some known participators are found, including *HLA-E* which is known to be up-regulated by cytomegalovirus infection, as well as *IL32* and *TNIP*, both upregulated and involved in host defense (Fig 2B). These genes were discovered before using cellular infection models (Fukushi et al. 1999; Ulbrecht et al. 2000; Kim et al. 2018). SMT is able to recapitulate results from these studies and provides more insights including more genes, under more natural setting in human, and in a spatially resolved way.

**Fig. 2.**
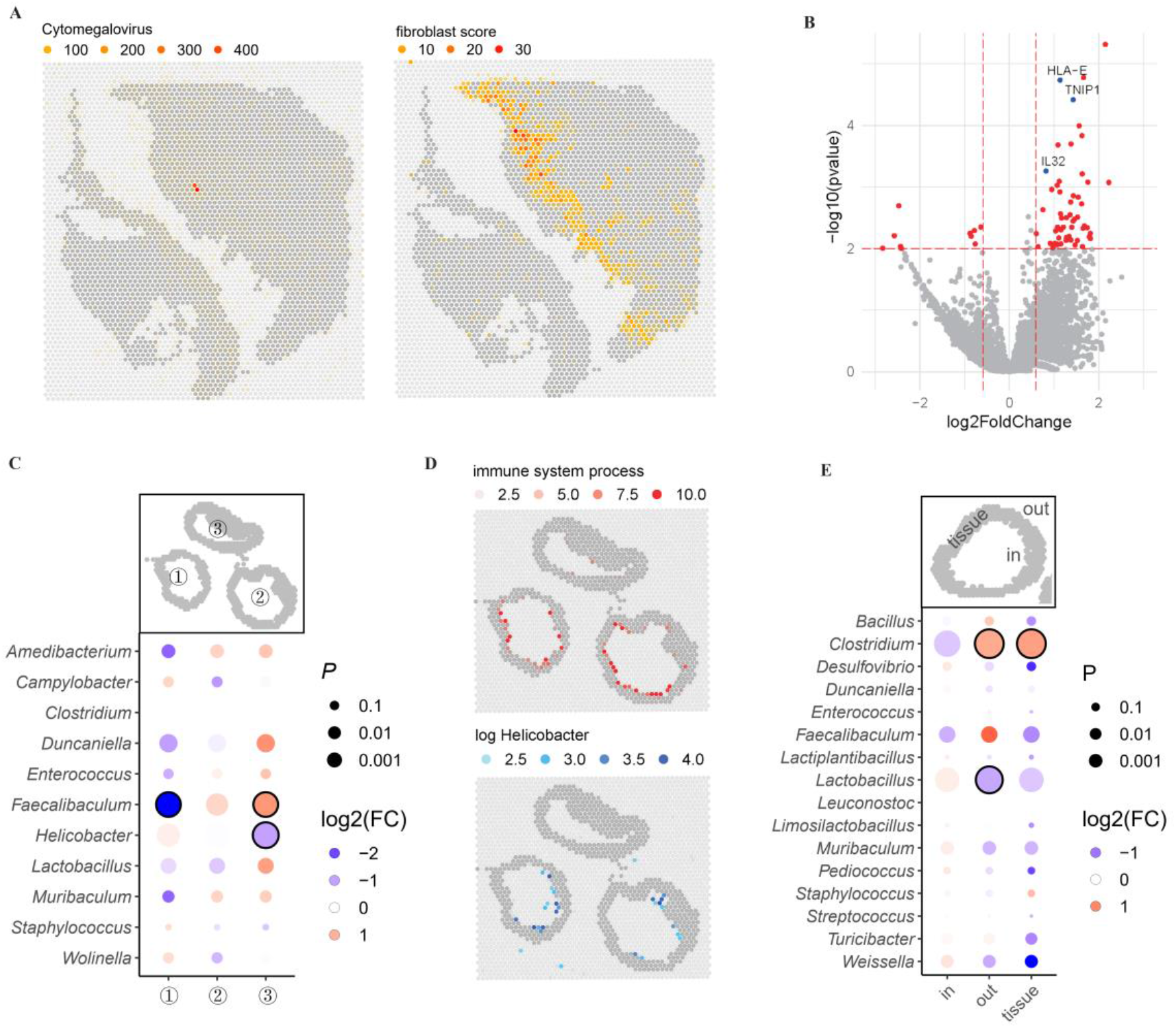
Host-microbe interactions revealed by SMT. **(A)** Spatial feature plot showing distribution of *Cytomegalovirus* in patient M1016’s section as detected by SMT (left) and distribution of fibroblasts as inferred by its marker gene score (right). **(B)** Volcano plot showing differentially expressed genes between infected and un-infected fibroblast spots, highly significant genes (p<0.01 and |log_2_FC|>0.5) were colored red, with several known ones labeled and highlighted in blue. **(C)** Dot plot showing the fold change and p value for microbial genera that were differentially distributed in the three sample murine samples as detected by Chi-square test, for mouse large intestine, those with p<10^−8^ and |log_2_FC| > 0.5 were circled. **(D)** Spatial feature plot showing module score of immune system process (top) and distribution of genus *Helicobacter* (left). **(E)** Dot plot showing fold change and p value for microbial genera that were differentially distributed in the three regions designated ‘in’, ‘tissue’ and ‘out’, by Chi-square test, for mouse small intestine, those with p<10^−8^ and |log_2_FC| > 0.5 were circled.

### SMT identifies highly immunogenic bacteria in murine intestine

The local cellular response to infection shown above is a good testament to SMT’s capability in investigating microbe-host interaction. Compared to other spatial method, FISH for example, SMT’s strength comes in its taxonomical resolution and systemic nature. For a proof of concept, we investigated the host-microbe interactions using their spatial correlations under homeostasis, using the murine dataset as it included multiple animals and covered complete cross-section of the intestines. A major challenge in this regard is the uncertainty or variation in the scale of interaction. Another technical issue is the sparse nature of the microbial signals and to a lesser degree, that of the host genes. To address both issues, a kernel smoothing function is applied to both host expression and microbial signals first (Methods, Fig S2A). The kernel width is adjustable and first degree neighbor was used by default. This way, small scale spatial interaction in the 100 um range, can be investigated using spatial correlation between the two smoothened signals. However, after multi-test correction, no interaction reached statistical significance. If we loosen the criteria, some suggestive interactions can be seem for the more abundant microbes like *Helicobacter* (Fig. S2B). Future development with improved capture efficiency and spatial resolution is required.

**Fig. S2. Related to Fig 2.**
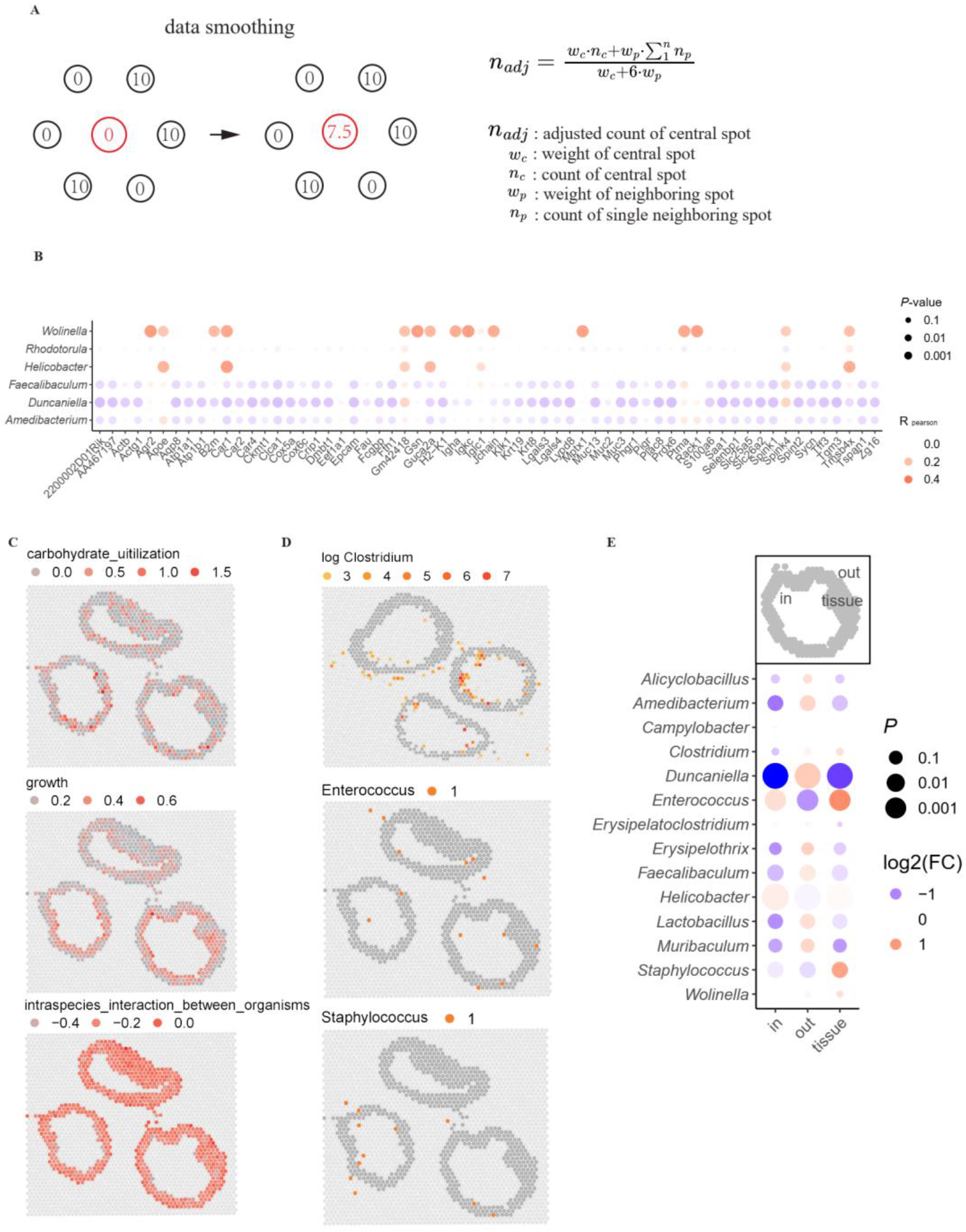
**(A)** Schema of kernel smoothing performed before microbe-host gene interaction detection. **(B)** Representative dot plot showing the spatial correlation between microbe and host genes in mouse large intestine, p values shown were before multi-test correction. **(C)** Representative spatial feature plot showing host pathways that did not show inter sample difference, in mouse large intestine. **(D)** Spatial feature plot for *Clostridium* in the small intestine, *Enterococcus* and *Staphylococcus* in the large intestine, showing their elevated concentration on tissue and further out, suggesting their trans-intestinal wall mobility. **(E)** Dot plot showing fold change and p value for microbial genera that were differentially distributed in the three regions designated ‘in’, ‘tissue’ and ‘out’, by Chi-square test, for mouse large intestine.

Zoom out to larger spatial scale interactions, we took a different approach wherein microbes and host pathways were evaluated on a per sample basis. There were three samples per window and sample level microbe abundances were compared. Specifically, microbial genera were tested for enrichment or depletion in particular samples, against background fluctuations, using Chi-square test and the microbes with per window UMI count larger than 5 were shown (Fig. 2C). On the other hand, host pathways were checked for differential enrichment amongst the different samples. Pathways were used instead of genes to reduce false positives. In the large intestine, *Helicobacter* is depleted in sample 3. Interestingly, out of the 31 top level GO pathway under Biological Processes, immune system process is also down-regulated in sample 3 (Fig. 2D, Fig. S2C). The correlation of *Helicobacter* with immune response was documented before and species in this genus was shown to be synergistic in tumor immune-therapy in mouse model (Overacre-Delgoffe et al. 2021). Our analysis suggest its high immunogenicity stands out from other bacteria even in seemingly homeostatic states. The difference between these homogenetic animals could be due to bona fide biological variation, and/or has to do with spatial variation in sample origin—whether they were more proximal or distal in the GI track and their proximity to lymphoid structures. A few other bacteria also showed inter-sample differences, *Faecalibaculum* in the large intestine for example (Fig. 2C). However their levels did now seem to closely track any top level pathways.

### SMT recapitulate the migration of *Clostridium* across intestinal wall

Bacteria species are generally confined in the intestinal lumen. However recent reports suggested certain bacteria, the most prominent being *Clostridium*, can migrate across the intestinal wall in inflammatory bowel disease and more strikingly, under homeostatic status (Ha et al. 2020). SMT provides a good opportunity to validate this observation and to investigate similar dynamics in a systematic way. To this end, we used the mouse intestinal dataset. For these slices, the spots under tissue, contained within the lumen or just outside the tissue were designated into regions ‘tissue’, ‘in’ or ‘out’ respectively. And for all bacteria with an overall frequency over 20, regional enrichment or depletion were tested using Chi-square test as before. In the small intestine, *Clostridium* standed out as having a significant enrichment on ‘tissue’ and ‘out’ (Fig. 2E), in agreement with previous report (Ha et al. 2020). In the large intestine, *Clostridium* presence was too low to yield any conclusion. Instead, *Enterococcus* and *Staphylococcus* were shown to have outward migratory capabilities with border-line significance (Fig. S2E). Nevertheless, we caution that these results are still preliminary and more SMT samples, preferably with better capture efficiency and resolution, in addition to orthogonal assays are needed to confirm and expand the discovery.

### Quantitative analysis of SMT’s sensitivity and recall using positive control

To quantitatively measure SMT’s sensitivity and recall, we carried out additional experiments using positive controls, and at the same time, tested a FISH inspired optimization for SMT. The experimental design is shown in Fig 3A. We utilized two windows on a Visium chip. Cultured control bacteria, including *Pseudomonas aeruginosa* and *Staphylococcus aureus*, were serial diluted and pipetted on three corners of a capture window, before tissue sections from the same patient were placed. The last corner was left out as blank control. Neighboring sections from the same tissue sample were used in the two windows. Then standard ST procedures were carried out. For one of the window, designated M1054B (patient M1054, replicate window B), an additional 15 minutes lysozyme treatment was performed after the permeabilization step (Methods). This step is commonly performed in FISH experiments and were shown to increase capture of diverse species with cell walls (Chassaing et al. 2014; Zou et al. 2018). After SMT data processing, clear circular shaped bacterial signal was evident at the two corners with the highest concentrations, with the next corner down in concentration still showing elevated bacteria load over the blank corner. The reason that there were two clear circular shape instead of three is likely the steep difference in bacteria concentration. The concentration was twenty-fold higher in one corner over the next and there was a 400 fold difference between the highest corner and the third one. Inevitably, some diffusion of molecules from the control bacteria did occur since they were actually placed outside the tissue (methods) and could diffuse away from the point of origin especially following the crevices at the outer edge of the tissue section Still, based on the strong contrast between the regions with positive control versus those immediately adjacent, one can expect for real tissue resident microbes, the diffusion effect should be less pronounced. We then calculated, for each corner with different control bacteria Colony Forming Unit (CFU), the ratio between UMI count and underlying CFU value. Factor in the diffusion effect, signal from one quadrant of a window was pooled together to represent control bacteria added in that corner. As shown in Fig 3B, the UMI to CFU ratio, at intermediate CFU level, is 0.55 UMI/CFU for *Staphylococcus* and 0.27 UMI/CFU for *Pseudomonas*. The trend lines stay linear between the highest two CFU levels and begin to level out due to diffusion effect at lower bacteria concentrations. These results suggest approximately one bacteria sequence captured per two to four bacteria cells. We caution that these estimates were crude since only two control bacteria were used and one of them, *Staphylococcus*, is Gram positive and have an additional lysis step applied (discussed later), which represented confounding factors. Microbial sequences other than the applied control bacteria were also quantified, and some “spillover” in taxonomy was evident as control bacteria sequences were called incorrectly, usually assigned to closely related organisms. These spillover signals can be identified as their per spot correlations with the control bacteria are abnormally high (Fig S3A). These spillover signals were calculated to be 8.13% for the *Staphylococcus* and 1.8% for *Pseudomonas*, representing a positive recall of over 90%.

**Fig. 3.**
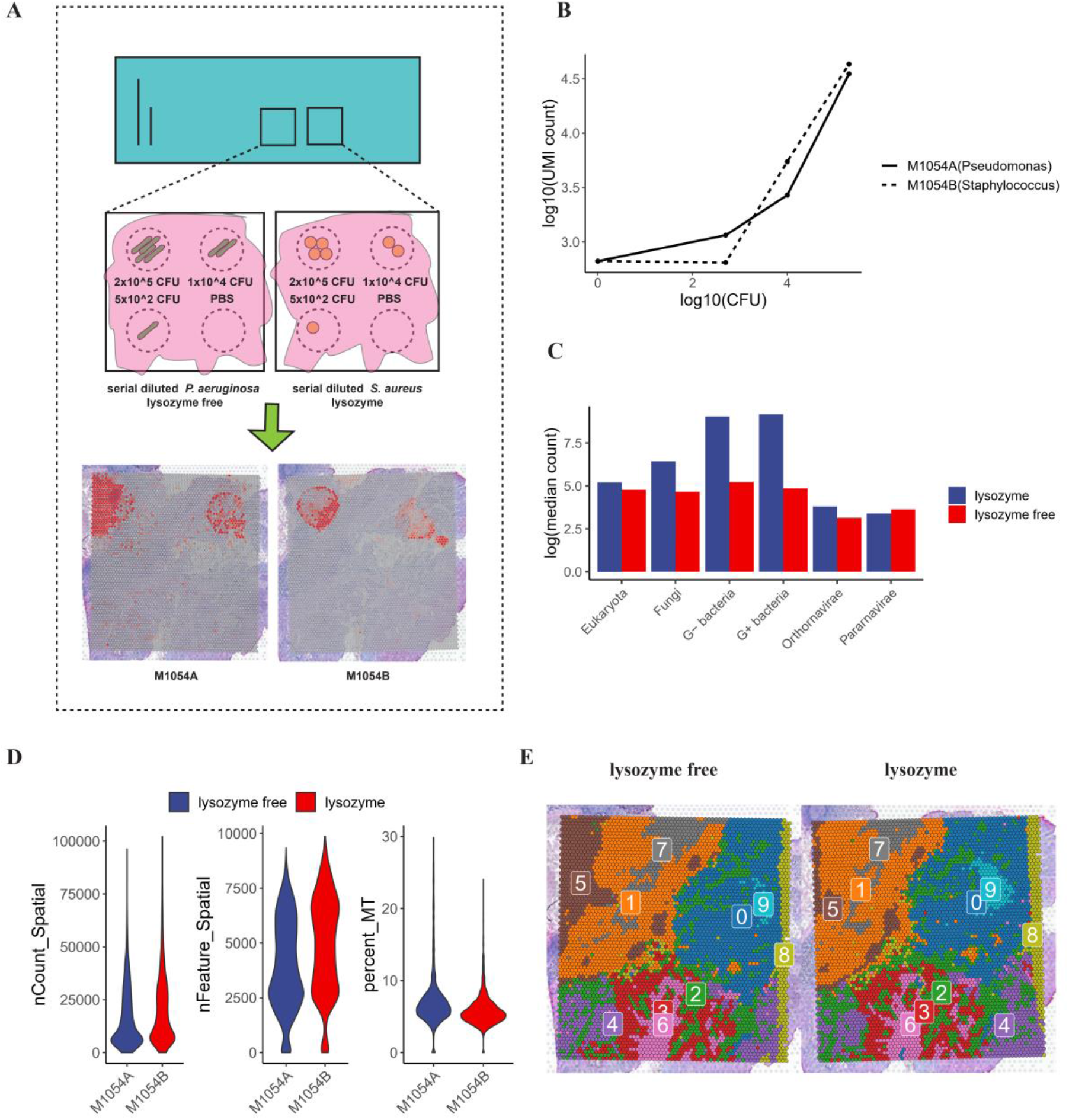
Capture efficiency and recall of SMT as measured by positive controls and evaluation of experimental optimization for SMT. **(A)** Study design of the positive control and optimization experiments, and shown at the bottom, spatial distribution of microbial sequences captured by SMT superimposed on tissue histology, showing clear circular patterns. **(B)** Control bacteria sequences captured by SMT at each corner (measured as total microbial UMI count, y-axis) against actual count of bacteria applied at that corner (measured as CFU, x-axis). **(C)** Bar plot showing microbial sequence capture with or without the additional lysozyme step. **(D)** Violin plots of host gene UMI count, gene count and mitochondria percentage (common measurements to QC single cell and ST data) with or without lysozyme treatment, no significant difference could be observed. **(E)** Spatial Dim Plots showing clustering of spatial spots using host expression data with or without lysozyme treatment, the similar pattern between the two conditions indicated the effect introduced by lysozyme treatment on host gene expression was minimal.

**Fig. S3. Related to Fig 3.**
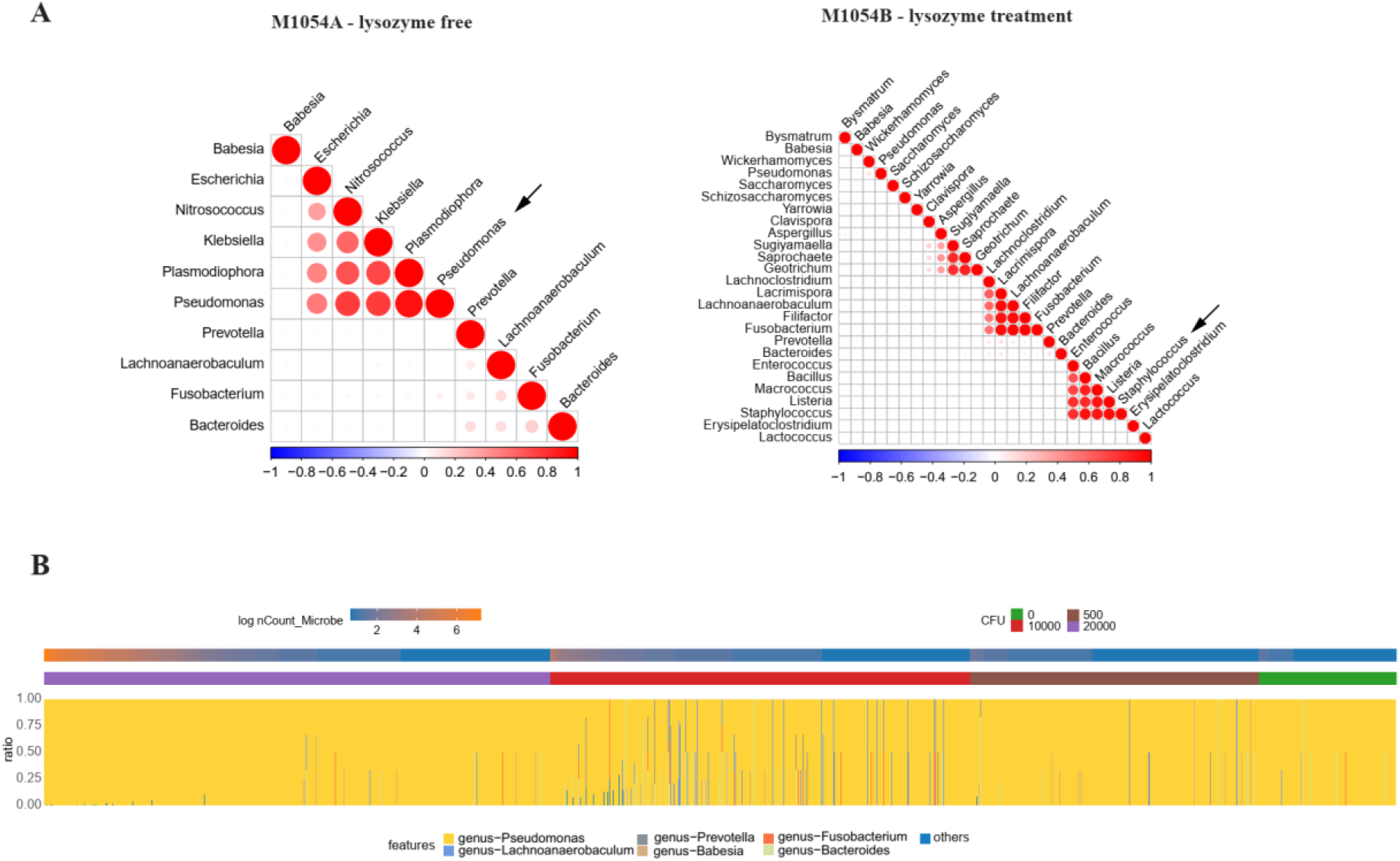
**(A)** Spatial correlation among bacteria genera as called by SMT, some spillover of the highly abundant control bacteria, *Pseudomonas* and *Staphylococcus*, did occur, manifesting as sequences assigned to closely related taxa. **(B)** Representative stacked bar plot showing microbial composition for individual spot in M1054A, their overall microbial sequence count and amount of control applied were also shown.

### Experimental optimization improves capture of microbes while maintaining host expression quality

With the control signals and their spillover filtered, the remaining microbial signals were treated as residential and used to evaluate the effect of additional lysis step in the experimental procedure. Between these two conditions, microbial signals from different kingdoms, with gram positive and negative bacteria separated, were compared. As expected, captured transcripts from gram positive and negative bacteria, prokaryotes and fungi are all significantly increased, with bacteria transcripts increased 4 folds on average (Fig 3C). At the same time, the per spot UMI count, gene count and mitochondria gene percentage, which are commonly used in quality check for single cell or spatial transcriptomic data, remained the same (Fig 3D). Furthermore, the spots from the two windows were clustered together and the resulting spatial pattern were nearly identical, some shift between the two sections notwithstanding (Fig 3E). In summary, the added lysozyme treatment greatly increased the microbial detection with no noticeable negative effect on the host gene expression.

## Discussion

Our work demonstrated the feasibility of Spatial Meta Transcriptomic sequencing and analysis, making possible for the first time simultaneous profiling by sequencing, of host tissue gene expression and microbial abundance in spatially resolved manner. Our analysis framework can extract spatial microbial abundance information from data generated by currently available commercial kits, effectively lifting suitable ST data to SMT status. We demonstrated that with careful experimental design and vigorous contaminant control and filtering, true signals can be discerned from background noises and that the biases were low. Moreover, comparisons with the FISH result suggested the current version of SMT method to be at similar levels of sensitivity. Nevertheless, we expect that in its current form, for significant findings in human, SMT likely will require large number of samples which can be an economical burden. Part of the reason has to do with technical issues like noise, limited resolutions etc., which will be ameliorated with future development. Another reason is shared among microbiome studies in that the microbiome is highly diverse and dynamic, especially in human. As an example, the two tumors used in this studies showed consistent difference in microbial load on their tumor tissue. This difference could be related to the fact that one of the tumor is Tertiary Lymphoid Structure positive and the other negative. Additional SMT dataset on the same type of tumor also supported the hypothesis the TLS is correlated with lower microbial load (data not shown), but has not reached statistical significance yet. Further SMT experiments or alternative assay aimed specifically at that hypothesis is needed.

The spatial transcriptomic technology that enabled SMT are rapidly evolving (Chen et al. 2021), reaching higher resolution and obtaining more sequences per area, translating to higher sensitivity. SMT too will benefit from these improvements. The current SMT framework can only generate a microbial abundance table, but in the foreseeable future, by simply optimizing microbial RNA extraction or by using specifically designed capture oligos, it may become possible to actually profile the microbial transcriptomes. Already, some next generation offerings of ST will replace poly T with random hexamer primer, which will benefit microbial sequences (personal communications with BGI group). Similarly, research is ongoing in our lab to generate higher resolution and more sensitive bacteria taxonomic information by adding an adapter oligo for Bacteria ribosomal RNA to the vanilla ST process. For tissues where microbes were abundant such as the intestines, SMT enables the systemic study of host and microbe interaction. In other more sterile tissues, SMT can reveal the consequence of microbial residence or intrusion, for example, help resolve how microbes travel to remote tumor site (as in non-intestinal solid cancers) and help settle the debate on whether certain body site is sterile or not, i.e. fetal organs (Kennedy et al. 2021; Mishra et al. 2021).

Through the past decade, microbiome sequencing has revolutionized the way microbe and host interactions are studied and most biological samples imaginable have had their microbiome sequenced. SMT is to bulk microbiome sequencing is analogous to single cell and ST sequencing to bulk RNA-Seq. Another interesting historical perspective was provided about a decade ago when RNA-Seq was just gaining popularity, researchers interested in infection dynamics coined the term ‘Dual RNA-seq’ to describe what was essentially meta-RNA-Seq used to investigate host and microbial expression simultaneously (Westermann et al. 2012). The work presented here paves the way for future development of SMT technology, which will see further increase sensitivity and taxonomical and spatial resolution. With their high throughput and systematic nature, SMT and technologies in the same vein will help address unmet challenges in the field of microbiome, immunology and beyond.

## Methods

### Human donors and murine samples processing

All human colon resection samples were obtained from colon cancer patients with written informed consent approved by the medical ethics committee of Ruijin hospital, Shanghai Jiao Tong University. Normal colon and tumor tissue were placed into conical tubes containing Roswell Park Memorial Institute (RPMI) media supplemented with 2% Fetal Bovine Serum (FBS) and placed on ice for transportation to the lab immediately, where they were cut into 1-2cm^3^ pieces, and embedded in Optimal Cutting Temperature (OCT) sectioning media (Thermo Scientific) by submersion in isopentane (2-methylbutane, Sigma Aldrich) pre-cooled to -80 °C in dry ice.

For murine samples, six-week-old male C57BL/6 mice were ordered from Shanghai SLAC Laboratory Animal Co. Ltd. All mice were maintained in the Specific Pathogen Free (SPF) mouse facilities at Fudan University. All experimental procedures described in the study were approved by the Institutional Animal Care and Use Committees (IACUC) of Fudan University. Feces and serum were collected and stored at -80 °C. Small intestine and colon tissue were washed in cold PBS and then embedded in OCT sectioning media (Thermo Scientific) by submersion in isopentane (2-methylbutane, Sigma Aldrich) pre-cooled to −80 °C in dry ice, and stored at −80 °C before Spatial Transcriptomic experiments.

### Spatial transcriptomics experiments(ST)and sequencing

Before ST experiment, cryosections were evaluated of their RNA integrity by taking 10 slides of 10 μM thickness, went through RNA extraction by Trizol (Invitrogen™,Cat#15596026) and analyzed by Agilent 2100. RNA integrity number (RIN) > 7 is considered qualified. Before using a new tissue for Visium Spatial Gene Expression libraries, the permeabilization time was optimized. The sections were fixed, stained and then underwent permeablization of different duration: 25 minutes, 18 minutes or 12 minutes. The permeabilization time that results in maximum fluorescence signal were selected. Once optimal conditions had been established, each section was cut at 10μM thickness and placed onto the Visium Spatial Gene Expression Slide (10X Genomics, Visium Spatial Gene Expression Slide & Reagent Kit, PN-1000187). Tissue sections were fixed, stained, and then scanned on a Leica DMI8 whole scanner at 10 X magnification. Then, for tissue permeabilization, the slides were placed in the Slide Cassette and Permeabilization Enzyme was added on the slide and incubated for the time selected in optimization step. After that, reverse transcription and second strand synthesis were on the slide, followed with cDNA quantification. The sequencing library construction was performed with Library Construction kit (10X Genomics, Cat#: PN-1000190). The cDNA was broken into fragments of about 200-300bp, and then the P7 and P5 adapter were added. Sequencing was performed using Illumina Novaseq 6000 sequencer with a sequencing depth targeting at least 50,000 reads per spot with pair-end 150 bp (PE150) reading strategy.

### ST data processing and analysis

Raw reads were aligned to the human transcriptome GRCh38-3.0.0 reference or mouse transcriptome mm10-3.0.0 reference using 10x Genomics SpaceRanger v.1.0.0 and exonic reads were used to produce mRNA count matrices for these samples. HE histology images were also aligned with mRNA capture spots using SpaceRanger.

Matrix file and HE image were then imported into R and analyzed using Seurat package (Version 4.0.1) (Butler et al. 2018). Downstream analysis was carried out following Seurat’s recommended procedure. Briefly, SpaceRanger output was imported into Seurat object, followed by normalization, scaling, variable feature identification, dimension reduction and clustering to find the general variation of the spots. Distribution of the major cell types were explored using the AddModuleScore function with cell type specific marker genes.

### SMT data analysis

An in-house pipeline was developed can take the bam file generated by SpaceRanger as input and output a microbial spatial abundance matrix. This matrix, together with the host expression one can be loaded into a Seurat object, and then spatially resolved microbe-host interaction can be explored using our custom codes. The SMT data analysis was described in the following steps.

#### 1) Spatial abundance matrix generation

Unmapped reads were extracted from the SpaceRanger bam file and filtered to remove low complexity sequences. The filtered reads were then aligned against nt database with blastn 2.10.1+. The blast output were formatted to preserve query ID, taxID, subject title, alignment length, e-value, identity, coverage as well as UMI and spatial barcode, which were extracted from the original bam file. Taxa levels from kingdom to species were called for individual mapped read using its corresponding taxID with the taxa table downloaded from NIH taxonomy portal. One UMI usually contain several reads. To make taxonomy calls for UMIs with conflicting read calls, we use the following strategy. If there is an over 80% majority, this majority call is used. Otherwise a last common ancestor of the conflicting calls is used. This way, a taxID-spatial barcode matrix could be generated.

#### 2) Removal of environmental contaminants

Before further analysis, microbial abundance matrices were denoised to remove likely contaminants that are distributed across the capture area in a random fashion. For this purpose, for every microbe, a number of random permutations of its placement were generated. The actual closeness of the spots containing the microbe was evaluated against the randomly generated distribution and a p value could be calculated. If spots containing a certain microbe were close to each other than could be expected by random chance, this usually imply that it is residential and its distribution follow certain structure. We optionally kept a known list of contaminants that can be used to filter the abundance matrix directly.

#### 3) Importing and using the abundance matrix

The abundance matrix and the host expression matrix were loaded into the same Seurat object for analysis. Specifically, a regular ST Seurat object was generated using the host expression first. The microbial abundance matrix was then processed into a phyloseq object (phyloseq 1.32.0) (McMurdie and Holmes 2013). This phyloseq object was then incorporated into the Seurat object as a separate assay. Custom codes were written that can generate abundance table at specified tax level on the fly and add that as additional assays. This way, the spatial distribution of the microbes could be investigated and visualized in the same way as host gene features.

#### 4) Spatial correlation between host genes and microbes

For spatial correlation analysis, expression data and microbial counts were first smoothened using a kernel method (Fig. S2A). With a kernel radius of one, the UMI count for the center spots was expanded to its nearest 6 spots with a weight of 0.25. Then correlation between microbes and genes could be carried out using spearman correlation or Kullback-Leibler divergence. When investigating interaction at larger spatial scale, for example, at the level of intestine sections or animals, the spatial spots were first designated into several regions and Chi-square test was used to find microbes that were significantly enriched or depleted in certain regions. In a similar fashion, enriched host expression features could be identified. Alternatively for top level host pathways (31 of them for GO Biological Process), highly variable ones can be identified by simple visualization.

### Comparison with Kraken2

To compare our Spatial Meta-Transcriptomic analysis pipeline with a commonly used tool Kraken2, data from M1016 was used for evaluation. Same unmapped reads were extracted from bam file from spaceranger output directory with samtools to generate a fasta file. After that, SMTA and Kraken2 were used separately for meta-transcriptomic profiling. The identity and coverage of SMTA were set to 80% and 60% respectively, while Kraken2 was run with default parameters.

We used the read per UMI metric as an indicator for the reliability of taxa assignment. As reads used for this purpose (unmapped to host) were enriched with low quality ones, and the low quality reads would generally have a low reads per UMI count while bona fide microbial UMI would have reads per UMI count similar to that of host gene UMIs (Table S1).

### Fluorescence in situ hybridization (FISH)

Optimal Cutting Temperature (O.C.T) embedded tissue slides were fixed with methyl alcohol at -20 °C. Slides were stained using the FISH kit (Guangzhou Exon Biotechnology Co.) following manufacturer’s protocol. Cy3 labelled Probes (EUB338-GCTGCCTCCCGTAGGAGT) were hybridized overnight at 37°C. Sections were washed at room temperature and 60°C in Washing Buffer I for 5 minutes, followed by two additional rounds of washing at 37°C for 5 minutes in Washing Buffer II. Staining was visualized with the Leica TCS SP8 confocal microscope at 20X/0.7 lens magnification and 63X/1.40 oil lens magnification. The images were analyzed using Imaris 9.7 Microscopy Imaging Software.

### Bulk RNA-Seq processing and analysis

OCT embedded tissue slices were used for RNA extraction with Trizol Reagent (#15596026, Invitrogen™) followed by library construction using Illumina Truseq Stranded Total RNA Kit (#20020596, Illumina™), and then went through RNA sequencing using Illumina Novaseq 6000 system (150-bp paired-end reads). The raw reads were aligned to the mouse reference genome (version mm10) and human reference genome (version hg38) using HISAT2 RNA-sequencing alignment software (Kim et al. 2019). The alignment files were processed to generate read counts for genes using SAMtools (Danecek et al. 2021) and HTSeq. Read counts were normalized and underwent differential analysis using R package DESeq2(Love et al. 2014). P values obtained from multiple tests were adjusted using the Benjamini-Hochberg correction.

### Experimental Optimization of SMT and related analysis

As ST experiment is not designed for microbial RNA capture, an experimental optimization following microbial FISH assay was tested. Our optimization followed the method used in (Chassaing et al. 2014). Specifically, after the tissue permeabilization step, a lysozyme treatment of 15 minutes was added (Beyotime Biotechnology Co., Cat#ST206), before the reverse transcription and second strand synthesis.

To compare the effects of lysozyme treatment, counts of resident microbes (positive controls removed), were normalized with global UMI counts and scaled to counts per 10^7^. Wilcoxon test was performed to compare the difference of microbial sequence abundance at kingdom level between lysozyme free and lysozyme treatment groups. Host expression qualities were evaluated by common metrics like feature per spot, UMI per spot and percentage of mitochondrial sequences, all using Seurat R package. Finally, host expressions under the two conditions were integrated and the spots clustered by the expression profiles and the two conditions were visualized by the Seurat SpatialDimPlot function.

### Using known bacteria as positive controls and related analysis

In order to validate and quantify SMT results, we used two control bacteria: a Gram positive *Staphylococcus aureus* and a Gram negative *Pseudomonas aeruginosa*. The bacteria were grown overnight for 16-18 hours at 37°C on a shaker, 250 rpm in LB broth supplemented with kanamycin. Bacteria were centrifuged at 5,000 rpm for 5 min, washed twice, and re-suspended in sterile PBS. The OD600 was measured to estimate bacterial density, and serial plating was performed on LB agar plates to quantify the use dose by counting colony forming units (CFU) after an overnight incubation at 37°C. The aim was to apply three different amounts of control bacteria at the three corners of a Visium capture window, 200,000 CFU, 10,000 CFU and 500 CFU respectively, and the last corner was left blank. Accordingly, the bacteria suspension was serial diluted and 0.5 uL of bacteria suspension was applied at the corners of capture area and let dry, before the ST experiments. During the ST process, the areas of application were generally covered by tissue.

For each Visium window with control bacteria added, the rectangular capture area was split evenly into four quadrants and each part was related to a particular gradient of added bacteria. In order to get rid of spillovers caused by ambiguous mapping, Spearman correlation analysis was performed, and those whose correlation coefficient with added control, i.e., *Pseudomonas* or *Staphylococcus*, larger than 0.5 were considered spillovers and were removed. The efficiency of control microbe detection was estimated by dividing total microbial UMI count from the second quadrant by its corresponding CFU count (10,000). This intermediate CFU was used as a plateauing of capture efficiency was observed at higher CFU.

## Data Access & Code Availability

The raw sequencing data for mouse was deposited in NCBI Short Read Archive with Bioproject ID PRJNA789453 (reviewer link https://dataview.ncbi.nlm.nih.gov/object/PRJNA789453?reviewer=dqhhb1320n1keabb58gvjps5g5). The raw data for human was deposited at the Genome Sequence Archive of China National Center for Bioinformation with accession ID HRA002678. Since this dataset is considered human genetic resources, raw data can be obtained by requesting and following the guidelines of Genome Sequence Archive for non-commercial use at https://ngdc.cncb.ac.cn/gsa-human/request/HRA002678. The guidance for making a data access request of GSA for humans can be downloaded from https://ngdc.cncb.ac.cn/gsa-human/document/GSA-Human_Request_Guide_for_Users_us.pdf. Code for the analysis in this manuscript is available at https://github.com/affinis/SMTa.

## Competing Interest Statment

The authors declare no competing interest.

## Acknowledgements

We would like to thank the Ruijin hospital of Shanghai for their help with tumor tissue preparation and specimen acquisition. We would also like to thank the sequencing core of Shanghai Institute of immunology for library construction and sequencing.

